# Functional Analysis of *cha* Genes Identifies ChaC as a Glutathione-Degrading Enzyme Rather Than a Sodium Transport Regulator

**DOI:** 10.64898/2026.05.15.725350

**Authors:** Hibiki Sawada, Naoko Ohkama-Ohtsu, Takehiro Ito

**Affiliations:** Graduate School of Agriculture, Tokyo University of Agriculture and Technology, Tokyo, Japan; Institute of Agriculture, Tokyo University of Agriculture and Technology, Tokyo, Japan; Research Unit of Nutrient Management, Advanced Research Center for One Welfare, Tokyo University of Agriculture and Technology, Tokyo, Japan; Graduate School of Agricultural and Life Sciences, The University of Tokyo, Tokyo, Japan

**Keywords:** *Escherichia coli*, γ-glutamylcyclotransferases, glutathione degradation, Na^+^/H^+^ antiporter, *cha* genes

## Abstract

Glutathione (GSH) is a tripeptide that plays essential roles in redox regulation and stress responses across organisms. In *Escherichia coli*, the GSH-specific γ-glutamyl cyclotransferase (ChaC) has been characterized biochemically, yet its physiological role remains unclear. Moreover, ChaC has been annotated as a regulator of the Na⁺/H⁺ antiporter ChaA based on its genomic association, although experimental evidence supporting this function is limited. In this study, we investigated whether *chaC* and its co-transcribed gene, *chaB*, are involved in sodium transport or GSH metabolism. Gene expression analyses revealed that *chaA*, *cha*B, and *chaC* are upregulated under salt stress. Functional analyses using deletion mutants showed that loss of *chaA* reduced salt tolerance, whereas deletion of *chaB* enhanced tolerance and decreased intracellular sodium levels. In contrast, deletion of *chaC* had no significant effect on salt tolerance or sodium accumulation. Overexpression of *cha* genes further indicated that *chaA*, but not *chaB* or *chaC*, contributed to salt tolerance. Importantly, overexpression of *chaC* significantly reduced intracellular GSH levels, whereas *chaB* overexpression had no effect. These results indicate that ChaC primarily functions in GSH degradation rather than in cation transport, and that ChaB does not participate in GSH metabolism. Our findings clarify the distinct physiological roles of ChaC and ChaB and provide new insight into bacterial physiology regarding GSH metabolism and ion transport in *E. coli*.

## Introduction

Living organisms contain diverse molecular components, among which glutathione (GSH; γ-glutamylcysteinylglycine) is an indispensable tripeptide that plays crucial roles in redox regulation, detoxification, oxidant scavenging, sulfur storage, and persulfide signaling [1–4]. GSH is synthesized via two ATP-dependent steps catalyzed by glutamate–cysteine ligase (GCL/γ-ECS) and GSH synthetase (GSS/GSH-S), and this pathway is essential for most organisms across kingdoms[5]. In *Escherichia coli*, Δ*gshA*, a mutant of GCL, is highly sensitive to oxidative stress [6], whereas Δ*gshB*, a mutant of GSS, shows increased sensitivity to cadmium [7]. In mice, the disruption of GCL or its catalytic subunit results in embryonic lethality [8–10]. In *Arabidopsis thaliana*, knockout of GCL and GSS causes embryonic and seedling lethality, respectively [11,12]. These observations highlight the conserved and fundamental importance of GSH across kingdoms.

Despite its importance, the GSH degradation pathway is also conserved among a wide variety of species [13]. In the cytosol, GSH-specific γ-glutamylcyclotransferases, ChaC, catalyze the cleavage of GSH to produce 5-oxoproline (5-OP) and Cys–Gly[14]. ChaC proteins are classified into two groups, ChaC1 and ChaC2 [13,14]. Whereas ChaC1 possesses high GSH-degrading activity and is found only in animals, ChaC2 shows low activity and is shared among bacteria, fungi, plants, and animals. In organisms possessing only ChaC2, including *E. coli*, ChaC2 is often referred to simply as ChaC. Despite its low activity, ChaC2 is constitutively expressed and is believed to slowly turn over GSH[13]. Nevertheless, its *in vivo* impact on GSH metabolism has not been fully elucidated.

This situation becomes more complex because ChaC is also suggested to have another function. *chaC* was initially identified in *E. coli* as a component of the ‘*cha* operon’ containing *chaA*, *chaB*, and *chaC* [15]. ChaA is a cation/H⁺ antiporter that was initially characterized as a Ca²⁺/H⁺ antiporter but was later identified as a Na⁺/H⁺ antiporter functioning at high pH [16–18]. In this context, ChaB and ChaC have been proposed as regulatory proteins of ChaA and remain annotated as cation transport regulators in databases and literature, although experimental support for these functions is limited.

These circumstances have led to two opposing interpretations of the roles of ChaC. Some studies interpret ChaC from the perspective of GSH catabolism, whereas others view it in the context of cation ion transport, particularly Na⁺. In *E.coli*, whereas the GSH degradation activity of ChaC has been demonstrated *in vitro* [13], its *in vivo* function has not been analyzed, and many databases and studies have annotated ChaC as a cation transport regulator to date [19–24]. In plants, Arabidopsis ChaC homologs (AtGGCTs) have been shown to degrade GSH, especially under sulfur deficiency and metal stress [25–27], whereas the rice ChaC homolog (OsARP) has been proposed to regulate Na⁺ transport into vacuoles [28–30]. Although recent studies have steadily accumulated evidence that ChaC acts as a GSH-degrading enzyme, its potential involvement in cation transport has not been examined, and the possibility of a mechanistic link between GSH degradation and cation transport cannot be ruled out.

Additionally, the function of ChaB also requires reinvestigation. Although ChaB has been studied under the assumption that it regulates ChaA, a functional relationship between ChaB and ChaA has not been experimentally verified to date [4,20,31–33]. Moreover, *chaB* is transcribed in the same orientation as *chaC* but in the opposite orientation as *chaA*, suggesting that the function of ChaB may be associated with GSH degradation by ChaC rather than with cation transport by ChaA.

In this study, we investigated whether the functions of ChaB and ChaC are associated with sodium transport via ChaA or with GSH degradation. We found that while *chaA* deletion reduced salt tolerance, *chaB* deletion enhanced it and lowered intracellular sodium levels. Meanwhile, *chaC* deletion did not affect salt tolerance or sodium levels. When overexpressed, whereas *chaA* increased salt tolerance, *chaB* or *chaC* had little or no impact on it. In contrast, *chaC* overexpression decreased GSH levels, whereas *chaB* overexpression did not affect them. These results indicate that the function of ChaC is associated with GSH degradation rather than ion transport via ChaA. In contrast, ChaB appears to play no role in GSH degradation and is instead potentially involved in sodium transport regulation. Our findings clarify the distinct physiological roles of ChaC and ChaB and provide new insight into bacterial physiology regarding GSH metabolism and ion transport in *E. coli*.

## Results

### The expression levels of *chaA*, *chaB*, and *chaC* are upregulated under salt stress

We first investigated the expression patterns of *cha* genes (Fig. 1). To examine whether *chaA*, *chaB*, and *chaC* respond to salt stress, reverse transcription quantitative PCR (RT-qPCR) was performed using cells grown in LBK medium containing 0, 200, or 400 mM NaCl at pH 8.8. The pH was adjusted to alkaline conditions because *chaA* is induced under alkaline conditions [18]. RT-qPCR analysis revealed that the transcript levels of *chaA*, *chaB*, and *chaC* were significantly higher under NaCl treatment than in the untreated control (Fig. 1B). At 200 mM NaCl, the transcript levels of *chaA*, *chaB*, and *chaC* increased by more than five-, seven-, and four-fold, respectively, and at 400 mM NaCl, by more than four-, nine-, and four-fold, respectively. To identify the transcriptional unit of *cha* genes, RT-PCR was performed using cells grown in LBK medium containing 0 mM NaCl at pH 8.8 (Fig. 1A and C). RT-PCR using primers spanning from the start of the *chaB* coding sequence (CDS) to the end of the *chaC* CDS yielded a band of the expected size, indicating that *chaB* and *chaC* were at least partially co-transcribed.

**Figure 1.**
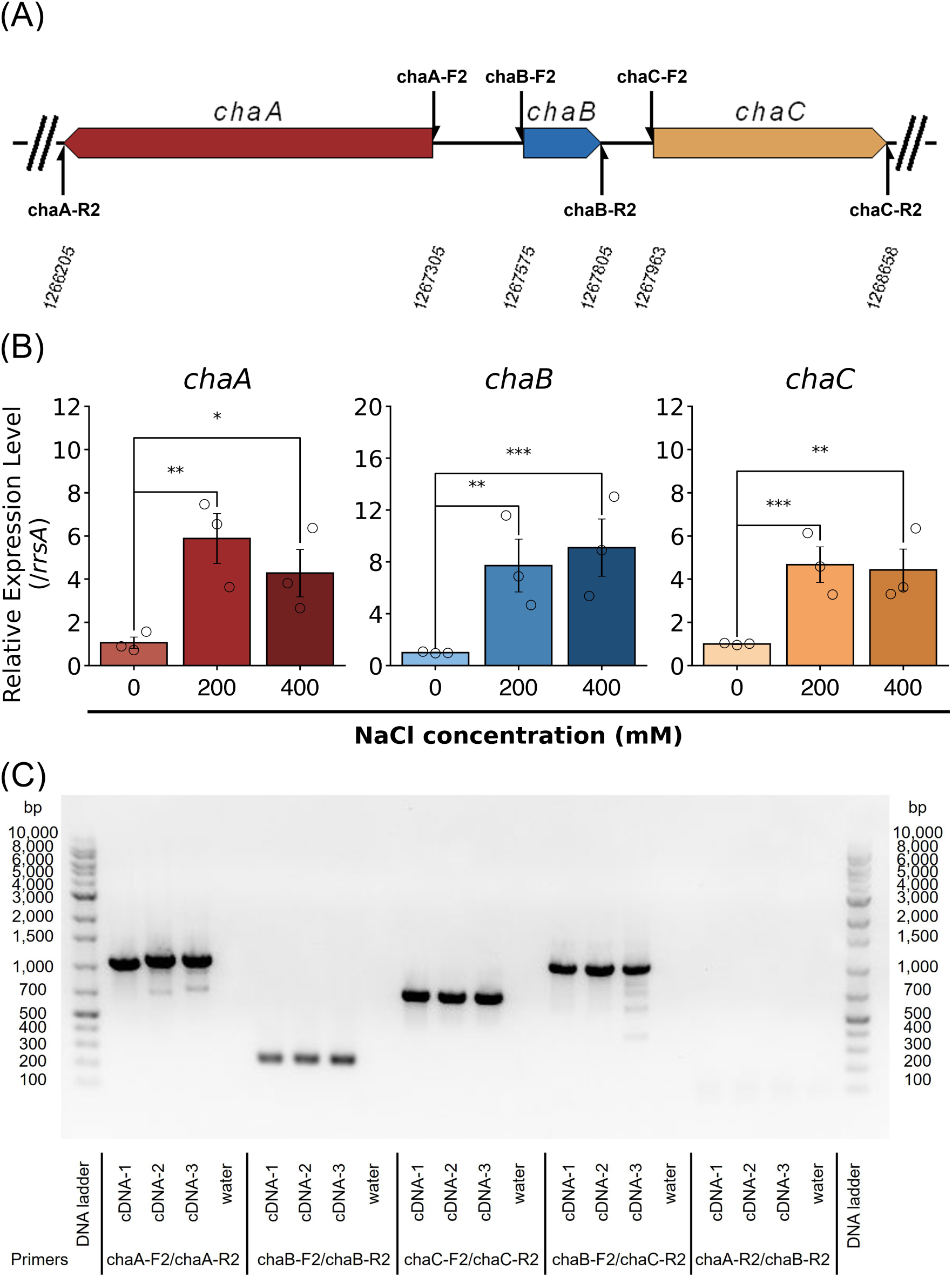
Transcriptional characteristics of *cha* genes. (A) Schematic diagram of the *cha* genes. The position and transcriptional direction of *chaA*, *chaB*, and *chaC* are shown. The numbers below the diagram indicate genomic positions of each gene. Half arrows indicate the binding sites of primers. (B) The relative expression levels of *chaA* (left), *chaB* (center), and *chaC* (right) in *Escherichia coli* exposed to 0, 200, or 400 mM NaCl at pH 8.8 at 37℃. The values are presented as mean ± standard error of the mean (SEM) of 2^^−ΔΔ*Ct*^ from three biological replicates. Δ*C_T_* values were used for statistical analysis. Statistical analyses were performed using one-way ANOVA followed by Dunnett’s multiple-comparison test. \**P* < 0.05, \*\**P* < 0.01, \*\*\**P* < 0.001. (C) RT-PCR was performed using cDNA synthesized from RNA isolated from cells grown in LBK medium containing 0 mM NaCl at pH 8.8 at 37℃. Each primer binding position is described in Fig. 1A. Primer pairs were designed within *chaA* (expected size: 482 bp), *chaB* (expected size: 1,100 bp), *chaC* (expected size: 695 bp), *chaB*-*chaC* (expected size: 1,083 bp), and *chaA*-*chaB* (expected size: 1,600 bp).

Furthermore, based on previous reports that *chaA* and *chaB* are regulated by the CpxAR two-component system [33], we analyzed the expression levels of *chaA*, *chaB*, and *chaC* in the Δ*cpxA* mutant in four independent experiments. While the previous study reported an increase in *chaA* and decrease in *chaB* mRNA levels in Δ*cpxA*, in our study, Δ*cpxA* showed a trend toward decreased *chaA* expression, while *chaB* and *chaC* expression varied among trials. (Fig. S1).

### Deletion of *chaB* enhances salt tolerance, whereas deletion of *chaC* has no effect on salt tolerance in the Δ*nhaA* background

To clarify the involvement of *chaA*, *chaB*, and *chaC* in salt tolerance, we examined the growth of the Δ*chaA*, Δ*chaB*, and Δc*haC* mutants under various salt concentrations at pH 8.8 (Fig. S2). However, no salt-sensitive phenotype was observed even in Δ*chaA*, making it difficult to evaluate the involvement of ChaB and ChaC in the regulation of ChaA. We reasoned that this might be due to the presence of the major sodium transporter NhaA, which could mask the effects of the *cha*A deletion, as indicated by Ohyama et al.[17]. Therefore, Δ*nhaA*Δ*chaA*, Δ*nhaA*Δ*chaB*, and Δ*nhaA*Δ*chaC* double mutants were created by P1 transduction and subjected to further analysis.

Δ*nhaA*Δ*chaA* were nonviable in the presence of NaCl at concentrations of 25 mM or higher, whereas Δ*nhaA*Δ*chaB* and Δ*nhaA*Δ*chaC* were viable in the presence of 0 – 400 mM NaCl (Fig. 2A and B). The area under the curve (AUC) of Δ*nhaA*Δ*chaB* was significantly lower than that of Δ*nhaA* at 0 mM NaCl, was comparable at 25 and 50 mM NaCl, and tended to be higher at 100, 200, and 400 mM NaCl, with a statistically significant difference at 200 mM NaCl. In contrast, no significant difference was observed between Δ*nhaA* and Δ*nhaA*Δ*chaC* at any concentration of NaCl (Fig. 2A and B).

**Figure 2.**
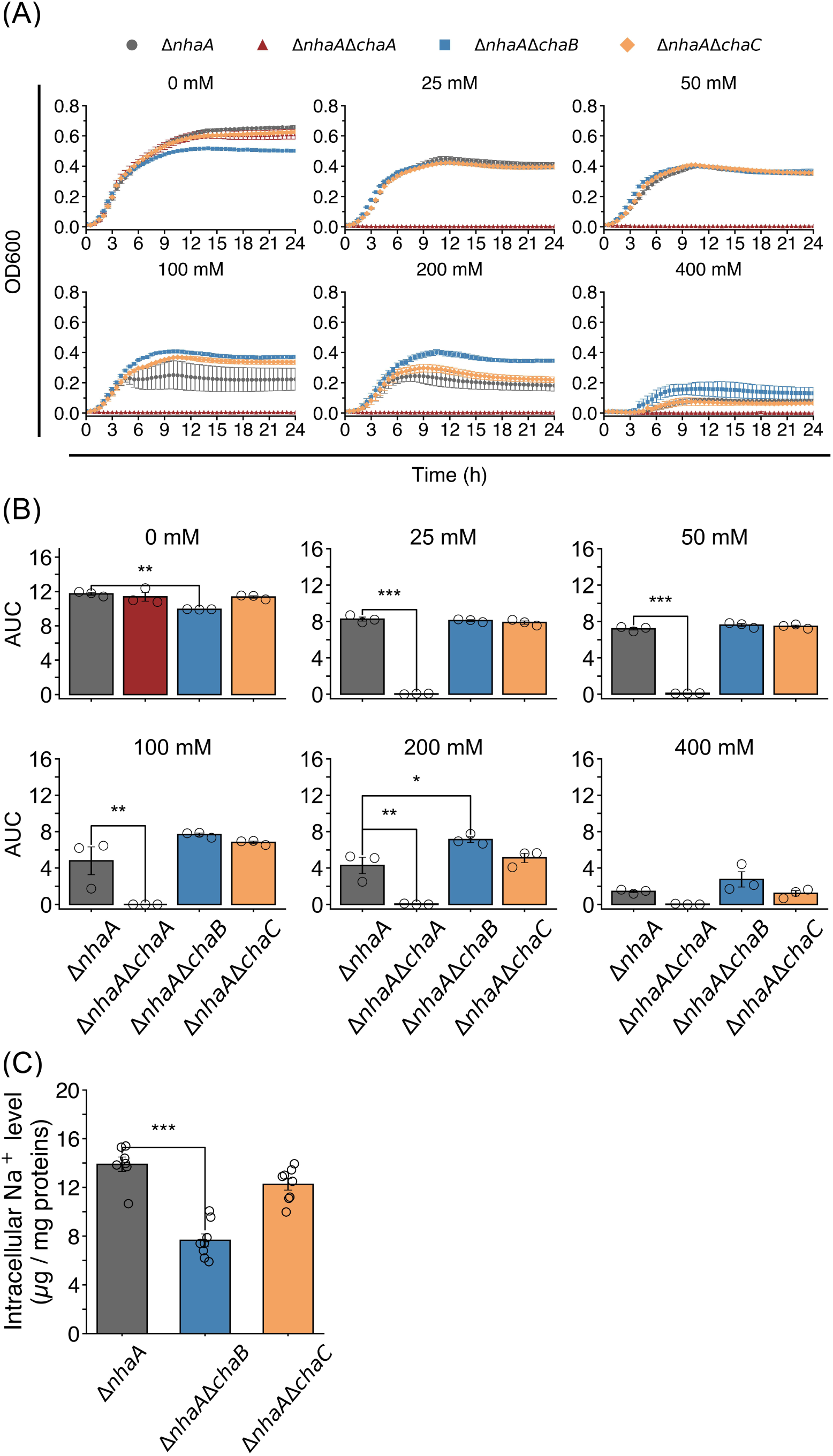
Effect of the *chaA*, *chaB*, or *chaC* deletion on salt sensitivity, growth and intracellular sodium levels in the Δ*nhaA* background at pH 8.8. (A) Growth curves of Δ*nhaA*, Δ*nhaA*Δ*chaA*, Δ*nhaA*Δ*chaB*, and Δ*nhaA*Δ*chaC* were generated by monitoring OD_600_ from 0 to 24 h using a microplate reader during incubation in LBK medium containing 0, 25, 50, 100, 200, or 400 mM NaCl at pH 8.8 at 37 °C. Raw OD_600_ values were corrected by subtracting blank measurements at each time point. (B) Area under the curve (AUC) values for Δ*nhaA*, Δ*nhaA*Δ*chaA*, Δ*nhaA*Δ*chaB*, and Δ*nhaA*Δ*chaC* were calculated from the growth curves shown in (A) over the 0 to 24 h interval using the R package *Growthcurver*. (C) Intracellular Na^+^ concentrations in Δ*nhaA*, Δ*nhaA*Δ*chaB*, and Δ*nhaA*Δ*chaC*. These mutants inoculated in LBK medium containing 200 mM NaCl at pH 8.8 at 37℃, cultured, and harvested at the logarithmic growth phase. The cells were disrupted by ultrasonication, and the NaCl concentration was measured by LAQUAtwin Na-11 analyzer (Horiba, Japan). For panel (A) and (B), values are presented as mean ± SEM from three biological replicates. For panel (C), values are presented as the mean ± SEM (n = 7–8). Statistical analyses were performed using one-way ANOVA followed by Dunnett’s multiple-comparison test. \**P* < 0.05, \*\**P* < 0.01, \*\*\**P* < 0.001.

To further evaluate whether changes in salt tolerance resulting from deletion of *chaB* or *chaC* in the Δ*nhaA* mutant are due to intracellular sodium levels, we measured sodium concentrations in the mutants at 200 mM NaCl. Compared with Δ*nhaA*, sodium concentration was significantly decreased in Δ*nhaA*Δ*chaB*, whereas Δ*nhaA*Δ*chaC* showed no significant difference (Fig. 2C).

Growth analysis was also performed at pH 7.0 (Fig. S3), where the contribution of ChaA is reportedly lower than at pH 8.8 [17,34]. Under these conditions, Δ*nhaA*Δ*chaA* was viable in the presence of 0 – 400 mM NaCl but exhibited lower OD_600_ values than Δ*nhaA* after approximately 9 h at 200 and 400 mM NaCl. In addition, the growth of Δ*nha*AΔ*chaB*, except at 100 mM, as well as that of Δ*nha*AΔ*chaC*, did not differ from that of Δ*nhaA* at pH 7.0.

### Leaky overexpression of *chaB*, *chaC* or *chaBC* does not largely alter salt tolerance

Subsequently, the overexpression lines of *chaA* (*chaA*_ox_), *chaB* (*chaB*_ox_), *chaC* (*chaC*_ox_), and *chaBC* (*chaBC*_ox_) were created by transforming the vector pBAD24-*chaA*, -*chaB*, - *chaC,* and -*chaBC* into the Δ*nhaA* strain, respectively (Fig. S4). *chaBC* represents a contiguous genomic sequence spanning from the beginning of the *chaB* CDS to the end of the *chaC* CDS. pBAD24 contains the arabinose-inducible pBAD promoter, which allows overexpression of the cloned gene in the presence of L-arabinose. However, elevated expression was observed even in the absence of arabinose induction (Fig. 3B) when grown at 200 mM NaCl at pH8.8. Relative to the empty vector control (EV), the expression level of *chaA* was more than eight-fold higher in *chaA*_ox_, that of *chaB* was more than five-fold higher in both *chaB*_ox_ and *chaBC*_ox_, and that of *chaC* was more than eleven-fold and sixty-fold higher in *chaC*_ox_ and *chaBC*_ox_, respectively.

**Figure 3.**
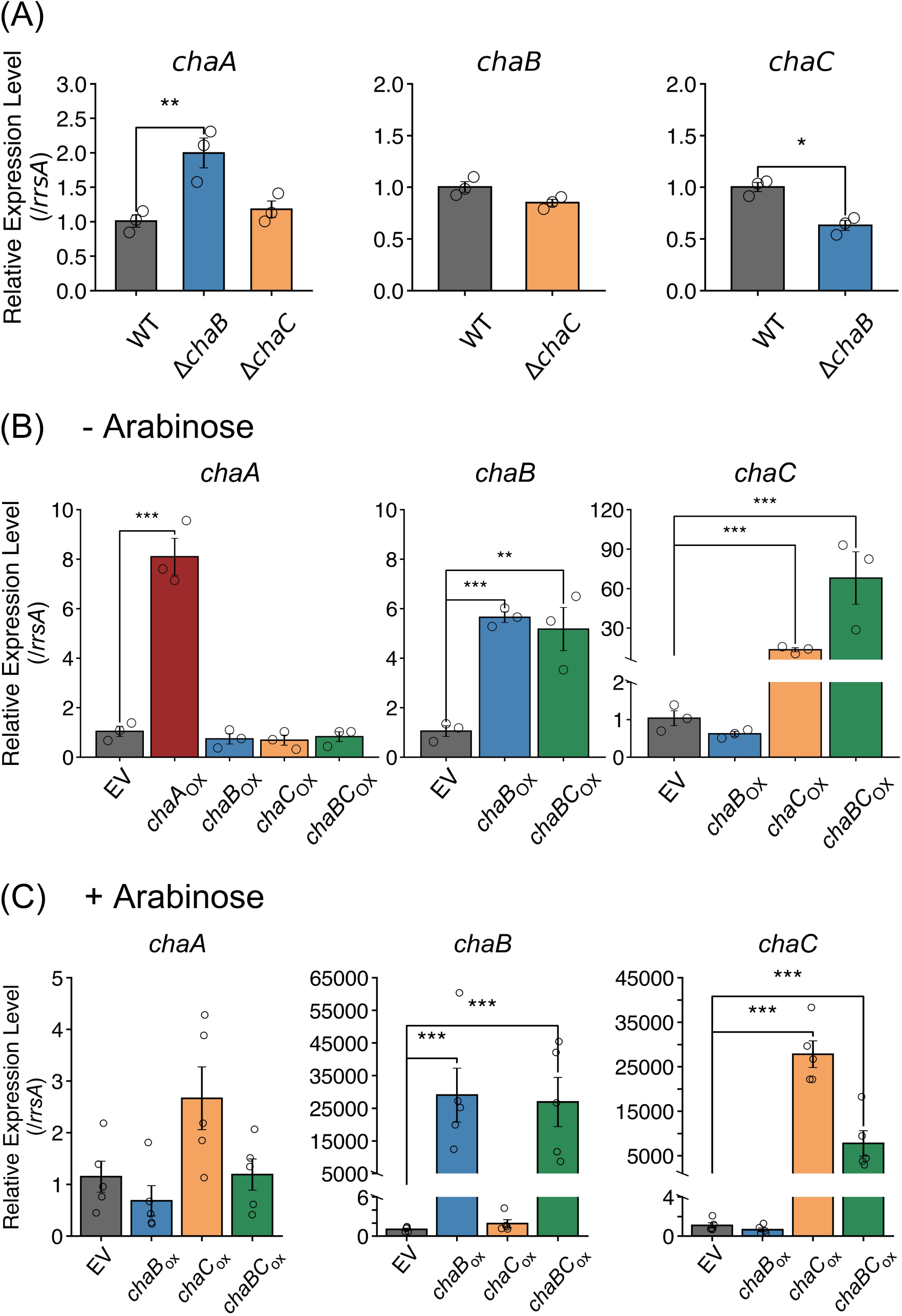
Impact of altered expressions of *chaB*, or *chaC* on the expression of *chaA*, *chaB*, and *chaC*. (A) The relative expression levels of *chaA* (left), *chaB* (center), *chaC* (right) in WT, Δ*chaB*, and Δ*chaC* exposed to 200 mM NaCl at pH 8.8. (B) The relative expression levels of *chaA* (left), *chaB* (center), and *chaC* (right) in *Escherichia coli* empty vector control (EV), *chaB*_ox_*, chaC*_ox_, and *chaBC*_ox_ exposed to 200 mM NaCl at pH 8.8 in the absence of L-arabinose. *chaBC* represents a contiguous genomic sequence spanning from the beginning of the *chaB* coding sequence to the end of the *chaC* coding sequence. (C) The relative expression levels of *chaA* (left), *chaB* (center), *chaC* (right) in EV, *chaB*_ox_*, chaC*_ox_, and *chaBC*_ox_ exposed to 200 mM NaCl at pH 8.8 in the presence of 0.02% L-arabinose. For panel (A) and (B), values are presented as the mean ± SEM of 2^^−ΔΔ*Ct*^ from three biological replicates. For panel (C), values are presented as the mean ± SEM of 2^^−ΔΔ*Ct*^ (n = 4–5). Δ*C_T_* values were used for statistical analysis. Statistical analysis was performed using Student’s *t*-test for panel A (center and right), and one-way ANOVA followed by Dunnett’s multiple-comparison test for panel A (left), B and C. \**P* < 0.05, \*\**P* < 0.01, \*\*\**P* < 0.001.

**Figure 4.**
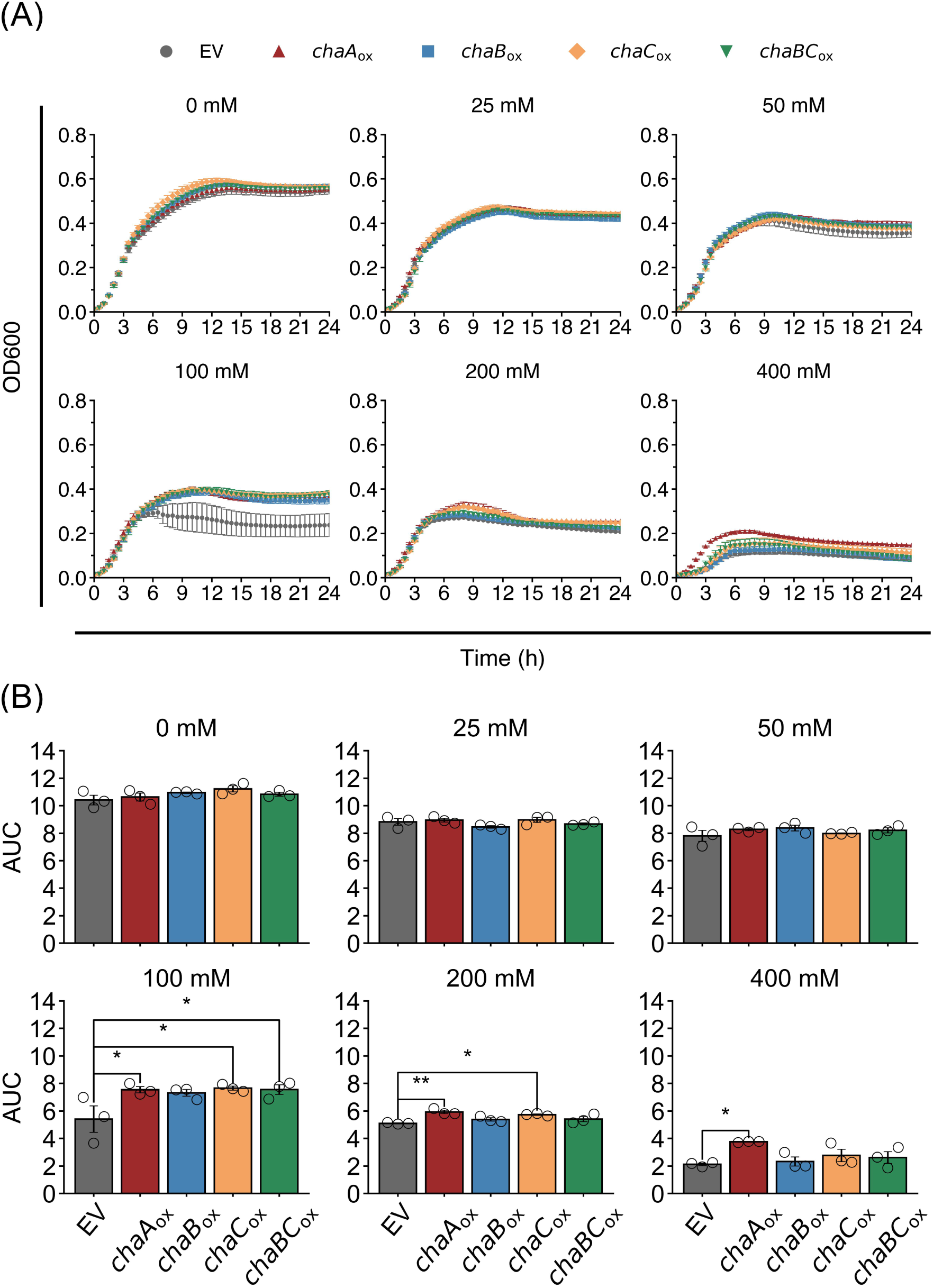
Effect of the *chaA, chaB*, or *chaC* overexpression on salt sensitivity and growth in Δ*nhaA* at pH 8.8 in the absence of L-arabinose. (A) Growth curves of empty vector control (EV), *chaA*-overexpression line (*chaA*_ox_), *chaB*-overexpression line (*chaB*_ox_), *chaC*-overexpression line (*chaC*_ox_), and *chaBC*-overexpression line (*chaBC*_ox_) were generated by monitoring OD_600_ from 0 to 24 h using a microplate reader during incubation in LBK medium containing 0, 25, 50, 100, 200, or 400 mM NaCl at pH 8.8 at 37 °C in the absence of L-arabinose. Raw OD_600_ values were corrected by subtracting blank measurements at each time point. (B) Area under the curve (AUC) values for EV, *chaA*_ox_, *chaB*_ox_, *chaC*_ox_, and *chaBC*_ox_ were calculated from the growth curves shown in (A) over the 0 to 24 h interval using the R package *Growthcurver*. For panel (A) and (B), values are presented as mean ± SEM from three biological replicates. Statistical analyses were performed using one-way ANOVA followed by Dunnett’s multiple-comparison test. \**P* < 0.05, \*\**P* < 0.01, \*\*\**P* < 0.001.

Thus, growth of the *chaA*_ox_, *chaB*_ox_, *chaC*_ox_, and *chaBC*_ox_ was examined under various salt concentrations at pH 8.8 without arabinose (Fig. 3B and C). Although no significant differences in AUC were detected in any of the strains under 0, 25, and 50 mM NaCl, *chaA*_ox_ showed higher AUC values at 100, 200 and 400 mM NaCl. Also, the AUC of *chaC*_ox_ was significantly higher at 100 and 200 mM NaCl, and that of *chaBC*_ox_ was significantly higher at 100 mM NaCl. However, the growth of EV was relatively poor, particularly at 100 mM NaCl, which may account for some observed significant differences. Thus, to confirm whether the overexpression of *chaC* affects the expression of *chaA*, RT-qPCR was performed using cells grown in LBK medium containing 200 mM NaCl at pH 8.8. However, no significant difference was observed (Fig. 3B). These results suggest that the enhanced salt tolerance of *chaC*_ox_ at 200 mM NaCl is unlikely to be attributed to *chaA*.

### GSH concentrations are decreased by the *chaC* overexpression but not by the *chaB* overexpression

Although ChaC was originally annotated as a regulator of ChaA [15], subsequent studies demonstrated that it functions as a GSH-specific γ-glutamyl cyclotransferase *in vitro* [13]. Despite this biochemical characterization, the physiological relevance of ChaC-mediated GSH degradation remains unclear in *E. coli* because no *in vivo* analyses using *chaC* knockout or overexpression lines have been reported to date. Furthermore, considering that *chaB* and *chaC* are co-transcribed (Fig. 1C), it is possible that ChaB also participates in the GSH degradation process.

To verify the involvement of ChaB and ChaC in GSH degradation *in vivo* and to investigate whether the altered salt tolerance in the mutants is attributed to GSH accumulation, we measured GSH levels in Δ*chaB* and Δ*chaC*. Compared with WT, GSH levels in Δ*chaB* and Δ*chaC* were not significantly different at pH 7.0 or pH 8.8. Similarly, no significant differences in GSH levels were observed in Δ*nhaA*Δ*chaB* and Δ*nhaA*Δ*chaC* at pH 8.8 (Fig. 5A). These results suggest that the deletion of *chaB* or *chaC* does not disrupt GSH homeostasis, possibly owing to the weak GSH degradation activity of ChaC, and also indicate that the increased salt tolerance of Δ*nhaA*Δ*chaB* compared to Δ*nhaA* (Fig. 2) is not attributed to GSH accumulation.

**Figure 5.**
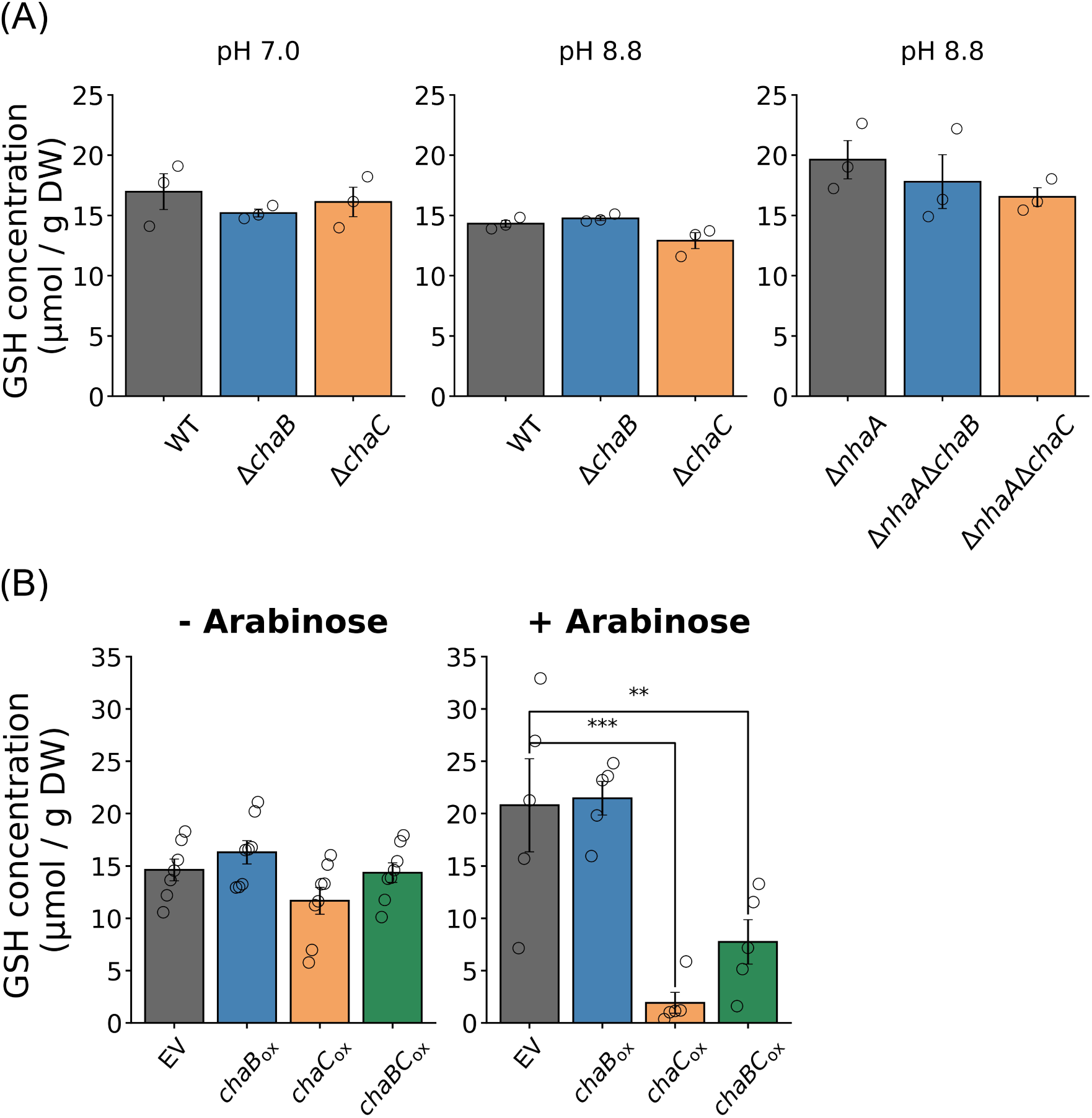
Effect of the deletion or overexpression of *chaB* and *chaC* on GSH concentrations. (A) GSH concentrations in WT, Δ*chaB*, and Δ*chaC* at pH 7.0 (left) and 8.8 (center), and in Δ*nhaA*, Δ*nhaA*Δ*chaB*, and Δ*nhaA*Δ*chaC* at pH 8.8 (right). These strains were inoculated in LBK medium containing 200 mM NaCl at pH 7.0 or 8.8 at 37℃, cultured, and harvested at the logarithmic growth phase. The cells were disrupted by ultrasonication, and the GSH concentration was measured by HPLC. (B) GSH concentrations in EV, *chaB*_ox_*, chaC*_ox_, and *chaBC*_ox_ in the absence (left) or in the presence (right) of L-arabinose. These strains inoculated in LBK medium containing 200 mM NaCl at pH 8.8 in the absence or presence of 0.02% L-arabinose, cultured, and harvested at the logarithmic growth phase. The cells were disrupted by ultrasonication, and the GSH concentration was measured by HPLC. For panel (A), values are presented as the mean ± SEM from three biological replicates. For panel (B), values are presented as the mean ± SEM (n = 4–8). Statistical analyses were performed using one-way ANOVA followed by Dunnett’s multiple-comparison test. \*\**P* < 0.01, \*\*\**P* < 0.001.

We also measured the GSH levels of EV, *chaB*_ox_, *chaC*_ox_, and *chaB*C_ox_ grown in LBK medium containing 200 mM NaCl at pH 8.8 without L-arabinose induction. Under these conditions, *chaB* and *chaC* expression levels are upregulated 5- to 70-fold (Fig. 3B). Nevertheless, no significant differences in GSH content were observed (Fig. 5B left).

To induce stronger expression, L-arabinose was added to the LBK medium to a final concentration of 0.02%. We first examined the expression levels of *chaC* and found that, relative to EV, they were not significantly different in *chaB*_ox_, whereas they were markedly higher in *chaC*_ox_ and *chaBC*_ox_, showing approximately 28,000- and 7,800-fold increases, respectively (Fig. 3C). Following RT-qPCR, we measured the GSH levels in EV, *chaB*_ox_, *chaC*_ox_ and *chaBC*_ox_ (Fig. 5B right). The GSH concentrations in *chaC*_ox_ and *chaBC*_ox_ were 9.1% and 37.5% of that of EV, respectively, whereas that of in *chaB*_ox_ was comparable to EV. These results indicate a negative correlation between *chaC* transcript levels and intracellular GSH concentrations and are consistent with the proposed role of ChaC in GSH degradation *in vivo*.

## Discussion

Although ChaC was originally annotated as a regulator of cation transporter ChaA [15], subsequent studies identified it as a GSH-specific γ-glutamylcyclotransferase [13]. Nevertheless, the initial interpretation as a ChaA regulator persists, likely because no conclusive data have been reported regarding the *in vivo* function of ChaC, leading to confused interpretations in some studies. Similarly, although ChaB has been regarded as a ChaA regulator based on its genomic proximity [15], limited evidence supports its involvement in salt tolerance, and no research has verified its potential role in ChaC-dependent GSH degradation. In this study, we show that ChaC functions in GSH degradation rather than in the regulation of ChaA-mediated sodium transport *in vivo*, and that ChaB may be involved in salt tolerance but not in GSH degradation.

### ChaC functions as a GSH degradation enzyme rather than as a cation transport regulator

Before verifying the roles of ChaC and ChaB in ChaA regulation, we first confirmed the function of ChaA as a Na^+^/H^+^ antiporter, as evidence for this is restricted to a few early reports. Our data suggested that, although NhaA plays a more prominent role in the wild type, ChaA is indispensable for salt tolerance in Δ*nhaA*, particularly under alkaline conditions (Fig. 2). These results are consistent with previous studies [16–18].

We subsequently investigated ChaC and found that it is unlikely to be involved in ChaA-mediated salt export. Unlike *chaA*, the deletion of *chaC* in the Δ*nhaA* background did not affect salt tolerance (Fig. 2A and B). Also, intracellular sodium concentrations in Δ*nhaA*Δ*chaC* were unchanged compared to Δ*nhaA* (Fig. 2C). Moreover, the expression levels of *chaA* in Δ*chaC* was unchanged compared with WT (Fig. 3A). These observations suggest that ChaC does not contribute to salt tolerance through the regulation of ChaA. To date, ChaC proteins have often been annotated as cation transport regulators across a wide variety of species, and, especially in rice, a ChaC homolog is estimated to confer salt tolerance by sodium transport regulation [28–30]. Our findings suggest that these interpretations may need to be reconsidered.

In contrast, our results indicate that ChaC functions as a γ-glutamylcyclotransferase for GSH *in vivo*, although its effect on GSH steady-state levels appears to be modest under normal conditions. In *chaC_ox_*, intracellular GSH levels were greatly reduced when its expression was induced with L-arabinose, whereas they were not significantly affected by basal expression (Fig. 3C). Additionally, the deletion of *chaC* did not alter GSH levels (Fig. 5A). These results support the GSH degradation ability of ChaC *in vivo*, whereas its influence on GSH turnover rate may be limited. Kaur et al.[13] reported that ChaC possesses only modest GSH degradation activity and is involved in a slow GSH turnover process, which is in consistent with our data. Despite its modest activity, ChaC proteins with similar kinetic parameters are well conserved across a broad range of species from bacteria to animals and plants[13]. The physiological role of ChaC proteins remains to be elucidated.

### The function of ChaB may be associated with ChaA rather than ChaC

In contrast to ChaC, our data suggest that ChaB may be associated with salt tolerance and the regulation of ChaA. Deletion of *chaB* in the Δ*nhaA* background resulted in increased salt tolerance and decreased intracellular sodium concentrations compared to Δ*nhaA* (Fig. 2). Furthermore, *chaA* expression was increased in Δ*chaB* (Fig. 3A). These results suggest that the *chaB* deletion enhanced salt tolerance by increasing *chaA* expression, implying that ChaB might act as a negative regulator of *chaA*.

However, this relationship can also be considered indirect or coincidental, as overexpression of *chaB* did not alter *chaA* expression levels, and both genes were upregulated under salt stress conditions (Fig. 1B and 3). In addition, a previous study has suggested that the functions of *chaA* and *chaB* are synergistic rather than antagonistic in antibiotic efflux, based on the increased sensitivity of Δ*chaA* and Δ*chaB* to antibiotic treatment [4]. Therefore, further studies are needed to conclude whether and how the functions of ChaA and ChaB are associated with each other.

In contrast to its role in salt tolerance, our data suggest that ChaB is not involved in ChaC-dependent GSH degradation. Although at least part of *chaB* and *chaC* are co-transcribed (Fig. 1C), neither the overexpression nor the deletion of *chaB* altered intracellular GSH levels (Fig. 5B). These results indicate that the physiological role of ChaB is distinct from that of ChaC.

## Conclusion

Although ChaC and ChaB were initially annotated as regulators of the cation transporter ChaA, the later identification of ChaC as a GSH-specific γ-glutamylcyclotransferase raised questions regarding their true functions. This study clarifies that ChaC is involved in GSH degradation rather than the regulation of ChaA-mediated sodium transport *in vivo*. In contrast, ChaB is unlikely to be associated with GSH degradation and instead is possibly involved in ChaA-mediated salt tolerance.

## Materials and Methods

### Growth conditions

Unless otherwise stated, *E.coli* were cultured at 37°C in LBK medium. LBK medium was prepared as LB broth without NaCl (10 g L⁻¹ tryptone and 5 g L⁻¹ yeast extract; pH was adjusted to 7.0 with KOH unless otherwise noted). When required, antibiotics were added at a final concentration of 100 μg mL⁻¹ for ampicillin and 50 μg mL⁻¹ for kanamycin.

### Competent cell preparation and heat shock transformation

Competent cells of each strain were prepared using the CaCl_2_ method. A 500 µl of pre-cultures were inoculated into 50 ml of LB medium and incubated at 37℃ with shaking at 180 rpm until reaching OD_600_ = 0.5. The cultures were transferred into a 50 ml centrifuge tube and centrifuged at 3,000 rpm for 5 min at 0°C. The supernatant was discarded and the cells were resuspended in 10 ml of 50 mM CaCl_2_. Then, the cells were centrifuged at 2,500 rpm for 3 min at 0°C. The supernatant was discarded and the cells were resuspended in 5 ml of 50 mM CaCl_2_ containing 15% glycerol. The cells were incubated on ice for 2h. After incubation, the cell suspension was aliquoted into 100 µl portions in microcentrifuge tubes and snap-frozen in liquid nitrogen. pCP20 (NovoPro, China) or pBAD24 (NIG, Japan) was transformed into competent cells by the heat shock method. To 100 µl of competent cells, 10 ng of plasmid was added and gently mixed. The mixture was incubated on ice for 30 min, heat-shocked at 42°C for 1 min, and then placed on ice for 3 min. Subsequently, 900 µl of SOC medium was added, and the cells were incubated at 30°C (for pCP20) or 37°C (for pBAD24) for 1 h.

### Establishment of single mutants and double mutants

All strains used in this study are listed in Table S1. WT (BW25113), *chaA::kan*, *chaB::kan*, *chaC::kan*, *nhaA::kan*, and *cpxA::kan*, which are parts of the Keio collection [35,36], were obtained from the National BioResource Project (NBRP, Japan). The kanamycin resistance gene cassette (*kan*) was removed from each strain using the pCP20 vector as described by Kuipers [13]. The resulting mutants were verified by colony PCR.

Double mutants (Δ*nhaA*Δ*chaA,* Δ*nhaA*Δ*chaB, and* Δ*nhaA*Δ*chaC*) were constructed by the P1 transduction following the protocol by Thomason et al. [37]. Bacteriophage P1 was provided by ATCC (USA). Briefly, P1 lysates prepared from the donor strains (*nhaA::kan*) were used to infect the recipient cells (Δ*chaA*, Δ*chaB*, *and* Δ*chaC*). The resulting transductants (*nhaA::kan*Δ*chaA, nhaA::kan*Δ*chaB, and nhaA::kan*Δ*chaC*) were selected on LB plates containing kanamycin. Then, Δ*nhaA*Δ*chaA,* Δ*nhaA*Δ*chaB, and* Δ*nhaA*Δ*chaC* were created by removing *kan* as described above. The resulting mutants were verified by colony PCR.

### Establishment of overexpression strains

The *chaA*, *chaB*, *chaC*, and *chaBC* fragments were amplified from *E. coli* BW25113 genomic DNA using the KOD DNA polymerase (Toyobo, Japan) and primers containing restriction sites for EcoRI and XbaI (Table S2). The PCR products were separated by agarose gel electrophoresis, and the DNA fragments of the expected size were excised and purified using the NucleoSpin Gel and PCR Clean-up Kit (Takara, Japan). The purified DNA fragments and the pBAD24 L-arabinose-inducible overexpression vector were double-digested with EcoRI and XbaI (New England Biolabs, USA). The reaction products were purified again and ligated using Ligation High Ver.2 (Takara, Japan). The ligation mixtures were transformed into *E. coli* DH5α competent cells by the heat shock method. Ampicillin-resistant colonies were screened on the selection plates, and the insertions were confirmed by colony PCR and Sanger sequencing (Eurofins Genomics, Japan) using primers listed in Table S2. The plasmids were collected using FastGene Plasmid Mini Kit (Nippon Genetics, Japan) and introduced into Δ*nhaA* by the heat-shock method. The cells were plated onto LB plates containing ampicillin and incubated at 37°C overnight to select transformants. Colonies were screened for the presence of the target plasmid insert by colony PCR using primers (Table S2) flanking the multiple cloning site, and the expected band size was confirmed.

### Quantitative RT-PCR

Each strain was inoculated into LBK medium. Cultures were incubated overnight at 37 °C with shaking at 180 rpm. After overnight cultivation, the optical density at 600 nm (OD₆₀₀) was measured using a GENESYS 30 Spectrophotometer (Thermo Fisher Scientific, USA). The cell suspension was diluted to OD₆₀₀ = 1.00 based on the spectrophotometric reading. 100 µL of the suspension (OD₆₀₀ = 1.00) was inoculated into 1.9 mL of fresh LBK medium (pH 8.8) containing 0, 200 or 400 mM NaCl (final concentration). Cultures were shaken at 37°C until reaching the logarithmic growth phase (OD₆₀₀ = 0.4–0.6). For the overexpression strains, L-arabinose was added at a final concentration of 0.02% (w/v). For RT-qPCR analysis, total RNA was isolated from bacterial cultures using the RNeasy Mini Kit (Qiagen, Germany; for Fig. 1, 3A and 3B) or RNAiso Plus (Takara, Japan; for Fig. S1, 3C and 3B). RNA concentration was determined using a NanoDrop Lite Plus spectrophotometer (Thermo Fisher Scientific, USA). Removal of genomic DNA and synthesis of cDNA were carried out using the PrimeScript RT reagent kit with gDNA Eraser (Takara, Japan). Primes used for RT-qPCR is listed in Table S2. The constitutively transcribed gene *rrsA* was used as a reference control to normalize the total RNA quantity of different samples. Differences between mRNA levels were calculated using the ΔΔ*C_T_* method. Δ*C_T_* values were used for statistical analysis.

### Growth analysis

A 10-µL aliquot of the OD₆₀₀ = 1.00 suspension was inoculated into 190 µL of LBK medium (pH 8.8) containing 0, 25, 50, 100, 200 or 400 mM NaCl (final concentration) in a 96-well microplate. The plate was incubated at 37°C in a Multiskan SkyHigh Microplate Spectrophotometer (Thermo Fisher Scientific, USA) with continuous cycles of 1-min shaking followed by a 1-min pause. OD₆₀₀ was recorded every 30 min for 24 h. AUC was calculated from each growth curve over the 0 to 24 h interval using the R package *Growthcurver* [38].

### GSH content determination

A 2.5-mL aliquot of the suspension (OD₆₀₀ = 1.00) was inoculated into 47.5 mL of LBK medium (pH 7.0 or 8.8) containing 200 mM NaCl (final concentration) in a 300-mL flask. For the overexpression strains, L-arabinose was added at a final concentration of 0.02% (w/v). Cultures were shaken at 37°C until reaching the logarithmic growth phase. The cultures were transferred to 50-mL centrifuge tubes and centrifuged at 5,000 × *g* for 10 min. The supernatant was discarded, and the pellets were freeze-dried. After freeze-drying, the dry weight of the cells was measured, and 10 mM HCl was added at 0.5-1.5 mL per mg dry weight. The suspension was sonicated on ice using an Ultrasonic Homogenizer THU-80 (AS ONE Corporation, Japan) to extract metabolites. Cell debris was removed by centrifugation (15,000 × *g* for 15 min at 4°C). The supernatant was used for GSH quantification by high-performance liquid chromatography (HPLC) after derivatization with monobromobimane [39,40]. The HPLC system (Shimadzu, Japan) consisted of a system controller (CBM-20A), two pumps (LC-20AD), a degasser (DGU-20A3), an autosampler (SIL-20A), a column oven (CTO-20 AC), and a fluorescence detector (RF-20A). A Shim-pack FC-ODS column (4.6 mm × 150 mm; Shimadzu, Japan) was used.

### Intracellular NaCl concentration determination

A 2.5-mL aliquot of the cell suspension (OD₆₀₀ = 1.00) was inoculated into 47.5 mL of LBK medium (pH 7.0 or 8.8) containing 200 mM NaCl (final concentration) in a 300-mL flask. Cultures were shaken at 37°C at 180 rpm until reaching the logarithmic growth phase. The cultures were transferred to 50-mL centrifuge tubes and centrifuged at 5,000 × *g* for 10 min, after which the supernatant was discarded. The cell pellets were washed four times by resuspending them in 1 mL of 0.3 M sucrose, centrifuging at 8,000 × *g* for 1 min at 4 °C, and removing the supernatant. After the final wash, the cells were resuspended in 1ml of Milli-Q water and completely disrupted by ultrasonication. The lysates were centrifuged at 20,000 × *g* for 10 min to remove cell debris. The supernatants were used for the determination of protein concentration and sodium ion levels. Protein concentrations were determined using the Bradford method, and sodium ion levels were measured using a LAQUAtwin Na-11 analyzer (Horiba, Japan).

## Data Availability

The data analyzed in the study are available within the article and its supplementary files.

## Open Access

Open access for this article was enabled through Read & Publish agreements between Portland Press and the University of Tokyo.

## CRediT Author Contribution

**Hibiki Sawada:** Conceptualization, Data curation, Formal analysis, Investigation, Visualization, Methodology, Writing - original draft. **Naoko Ohkama-Ohtsu:** Conceptualization, Resources, Supervision, Methodology, Writing - review & editing. **Takehiro Ito:** Conceptualization, Resources, Supervision, Funding acquisition, Methodology, Project administration, Writing - review & editing.

## Acknowledgements

We are grateful to the National BioResource Project (NIG, Japan) for providing the *E. coli* strains from the Keio collection.

## References

[1] Vignane T, Filipovic MR. Emerging Chemical Biology of Protein Persulfidation. Antioxid Redox Signal 2023;39:19–39. 10.1089/ARS.2023.0352;ISSUE:ISSUE:DOI.

[2] Forman HJ, Zhang H, Rinna A. Glutathione: Overview of its protective roles, measurement, and biosynthesis. Mol Aspects Med 2009;30:1–12. 10.1016/J.MAM.2008.08.006.

[3] Ito T, Ohkama-Ohtsu N. Degradation of glutathione and glutathione conjugates in plants. J Exp Bot 2023;74:3313–27. 10.1093/JXB/ERAD018.

[4] Wan Y, Wang M, Chan EWC, Chen S. Membrane Transporters of the Major Facilitator Superfamily Are Essential for Long-Term Maintenance of Phenotypic Tolerance to Multiple Antibiotics in E. coli. Microbiol Spectr 2021;9. 10.1128/SPECTRUM.01846-21;WGROUP:STRING:PUBLICATION.

[5] Noctor G, Gomez L, Vanacker H, Foyer CH. Interactions between biosynthesis, compartmentation and transport in the control of glutathione homeostasis and signalling. J Exp Bot 2002;53:1283–304. 10.1093/JEXBOT/53.372.1283.

[6] Oktyabrskii ON, Muzyka NG, Ushakov VY, Smirnova G V. The role of thiol redox systems in the peroxide stress response of Escherichia coli. Microbiology 2007 76:6 2007;76:669–75. 10.1134/S0026261707060045.

[7] Helbig K, Bleuel C, Krauss GJ, Nies DH. Glutathione and transition-metal homeostasis in Escherichia coli. J Bacteriol 2008;190:5431–8. 10.1128/JB.00271-08;SUBPAGE:STRING:FULL.

[8] Shi ZZ, Osei-Frimpong J, Kala G, Kala S V., Barrios RJ, Habib GM, et al. Glutathione synthesis is essential for mouse development but not for cell growth in culture. Proc Natl Acad Sci U S A 2000;97:5101–6. 10.1073/PNAS.97.10.5101;SUBPAGE:STRING:FULL.

[9] Winkler A, Njålsson R, Carlsson K, Elgadi A, Rozell B, Abraham L, et al. Glutathione is essential for early embryogenesis – Analysis of a glutathione synthetase knockout mouse. Biochem Biophys Res Commun 2011;412:121–6. 10.1016/J.BBRC.2011.07.056.

[10] Dalton TP, Dieter MZ, Yang Y, Shertzer HG, Nebert DW. Knockout of the Mouse Glutamate Cysteine Ligase Catalytic Subunit (Gclc) Gene: Embryonic Lethal When Homozygous, and Proposed Model for Moderate Glutathione Deficiency When Heterozygous. Biochem Biophys Res Commun 2000;279:324–9. 10.1006/BBRC.2000.3930.

[11] Pasternak M, Lim B, Wirtz M, Hell R, Cobbett CS, Meyer AJ. Restricting glutathione biosynthesis to the cytosol is sufficient for normal plant development. Plant Journal 2008;53:999–1012. 10.1111/J.1365-313X.2007.03389.X;PAGE:STRING:ARTICLE/CHAPTER.

[12] Cairns NG, Pasternak M, Wachter A, Cobbett CS, Meyer AJ. Maturation of Arabidopsis Seeds Is Dependent on Glutathione Biosynthesis within the Embryo. Plant Physiol 2006;141:446–55. 10.1104/PP.106.077982.

[13] Kaur A, Gautam R, Srivastava R, Chandel A, Kumar A, Karthikeyan S, et al. ChaC2, an Enzyme for Slow Turnover of Cytosolic Glutathione. Journal of Biological Chemistry 2017;292:638–51. 10.1074/JBC.M116.727479.

[14] Kumar A, Tikoo S, Maity S, Sengupta S, Sengupta S, Kaur A, et al. Mammalian proapoptotic factor ChaC1 and its homologues function as γ-glutamyl cyclotransferases acting specifically on glutathione. EMBO Rep 2012;13:1095–101. 10.1038/EMBOR.2012.156/FIGURES/4.

[15] Oshima T, Aiba H, Baba T, Fujita K, Hayashi K, Honjo A, et al. A 718-kb DNA Sequence of the Escherichia coli K-12 Genome Corresponding to the 12.7–28.0 min Region on the Linkage Map. DNA Research 1996;3:137–55. 10.1093/DNARES/3.3.137.

[16] Ivey DM, Guffanti AA, Zemsky J, Pinner E, Karpel R, Padan E, et al. Cloning and characterization of a putative Ca2+/H+ antiporter gene from escherichia coli upon functional complementation of Na+/H+ antiporter-deficient strains by the overexpressed gene. Journal of Biological Chemistry 1993;268:11296–303. 10.1016/s0021-9258(18)82124-x.

[17] Ohyama T, Igarashi K, Kobayashi H. Physiological role of the chaA gene in sodium and calcium circulations at a high pH in Escherichia coli. J Bacteriol 1994;176:4311–5. 10.1128/JB.176.14.4311-4315.1994;CTYPE:STRING:JOURNAL.

[18] Shijuku T, Yamashino T, Ohashi H, Saito H, Kakegawa T, Ohta M, et al. Expression of chaA, a sodium ion extrusion system of Escherichia coli, is regulated by osmolarity and pH. Biochimica et Biophysica Acta (BBA) - Bioenergetics 2002;1556:142–8. 10.1016/S0005-2728(02)00345-6.

[19] Hua Q, Yang C, Oshima T, Mori H, Shimizu K. Analysis of Gene Expression in Escherichia coli in Response to Changes of Growth-Limiting Nutrient in Chemostat Cultures. Appl Environ Microbiol 2004;70:2354–66. 10.1128/AEM.70.4.2354-2366.2004;ISSUE:ISSUE:DOI.

[20] Hayes ET, Wilks JC, Sanfilippo P, Yohannes E, Tate DP, Jones BD, et al. Oxygen limitation modulates pH regulation of catabolism and hydrogenases, multidrug transporters, and envelope composition in Escherichia coli K-12. BMC Microbiology 2006 6:1 2006;6:89-. 10.1186/1471-2180-6-89.

[21] White-Ziegler CA, Um S, Pérez NM, Berns AL, Malhowski AJ, Young S. Low temperature (23 °C) increases expression of biofilm-, cold-shock- and RpoS-dependent genes in Escherichia coli K-12. Microbiology (N Y) 2008;154:148–66. 10.1099/MIC.0.2007/012021-0.

[22] Vanaja SK, Springman AC, Besser TE, Whittam TS, Manning SD. Differential expression of virulence and stress fitness genes between Escherichia coli O157:H7 strains with clinical or bovine-biased genotypes. Appl Environ Microbiol 2010;76:60–8. 10.1128/AEM.01666-09;WEBSITE:WEBSITE:ASMJ;JOURNAL:JOURNAL:AM;ISSUE:ISSUE:DOI.

[23] Phadtare S. Escherichia coli cold-shock gene profiles in response to over-expression/deletion of CsdA, RNase R and PNPase and relevance to low-temperature RNA metabolism. Genes to Cells 2012;17:850–74. 10.1111/GTC.12002;REQUESTEDJOURNAL:JOURNAL:13652443;CTYPE:STRING:JOURNAL.

[24] Wang Z, Xue T, Hu D, Ma Y. A Novel Butanol Tolerance-Promoting Function of the Transcription Factor Rob in Escherichia coli. Front Bioeng Biotechnol 2020;8:524198. 10.3389/FBIOE.2020.524198/TEXT.

[25] Joshi NC, Meyer AJ, Bangash SAK, Zheng ZL, Leustek T. Arabidopsis γ-glutamylcyclotransferase affects glutathione content and root system architecture during sulfur starvation. New Phytologist 2019;221:1387–97. 10.1111/NPH.15466;ISSUE:ISSUE:DOI.

[26] Paulose B, Chhikara S, Coomey J, Jung H Il, Vatamaniuk O, Dhankher OP. A γ-Glutamyl Cyclotransferase Protects Arabidopsis Plants from Heavy Metal Toxicity by Recycling Glutamate to Maintain Glutathione Homeostasis. Plant Cell 2013;25:4580–95. 10.1105/TPC.113.111815.

[27] Kumar S, Kaur A, Chattopadhyay B, Bachhawat AK. Defining the cytosolic pathway of glutathione degradation in Arabidopsis thaliana: role of the ChaC/GCG family of γ-glutamyl cyclotransferases as glutathione-degrading enzymes and AtLAP1 as the Cys-Gly peptidase. Biochemical Journal 2015;468:73–85. 10.1042/BJ20141154.

[28] Qi Y, Yamauchi Y, Ling J, Kawano N, Li D, Tanaka K. The submergence-induced gene OsCTP in rice (Oryza sativa L.) is similar to Escherichia coli cation transport protein ChaC. Plant Science 2005;168:15–22. 10.1016/J.PLANTSCI.2004.07.004.

[29] Uddin MI, Qi Y, Yamada S, Shibuya I, Deng XP, Kwak SS, et al. Overexpression of a New Rice Vacuolar Antiporter Regulating Protein OsARP Improves Salt Tolerance in Tobacco. Plant Cell Physiol 2008;49:880–90. 10.1093/PCP/PCN062.

[30] Uddin MI, Kihara M, Yin L, Perveen MF, Tanaka K. Expression and subcellular localization of antiporter regulating protein OsARP in rice induced by submergence, salt and drought stresses. Afr J Biotechnol 2012;11:12849–55. 10.5897/AJB11.2224.

[31] Osborne MJ, Siddiqui N, Iannuzzi P, Gehring K. The solution structure of ChaB, a putative membrane ion antiporter regulator from Escherichia coli. BMC Structural Biology 2004 4:1 2004;4:9-. 10.1186/1472-6807-4-9.

[32] Sukumaran A, Pladwig S, Geddes-McAlister J. Zinc limitation in Klebsiella pneumoniae profiled by quantitative proteomics influences transcriptional regulation and cation transporter-associated capsule production. BMC Microbiology 2021 21:1 2021;21:43-. 10.1186/S12866-021-02091-8.

[33] Zhao Z, Xu Y, Jiang B, Qi Q, Tang Y-J, Xian M, et al. Systematic Identification of CpxRA-Regulated Genes and Their Roles in Escherichia coli Stress Response. MSystems 2022;7. 10.1128/MSYSTEMS.00419-22;PAGE:STRING:ARTICLE/CHAPTER.

[34] Sakuma T, Yamada N, Saito H, Kakegawa T, Kobayashi H. pH dependence of the function of sodium ion extrusion systems in Escherichia coli. Biochimica et Biophysica Acta (BBA) - Bioenergetics 1998;1363:231–7. 10.1016/S0005-2728(97)00102-3.

[35] Baba T, Ara T, Hasegawa M, Takai Y, Okumura Y, Baba M, et al. Construction of Escherichia coli K-12 in-frame, single-gene knockout mutants: the Keio collection. Molecular Systems Biology 2006 2:1 2006;2:MSB4100050-. 10.1038/MSB4100050.

[36] Yamamoto N, Nakahigashi K, Nakamichi T, Yoshino M, Takai Y, Touda Y, et al. Update on the Keio collection of Escherichia coli single-gene deletion mutants. Molecular Systems Biology 2009 5:1 2009;5:MSB200992-. 10.1038/MSB.2009.92.

[37] Thomason LC, Costantino N, Court DL. E. coli Genome Manipulation by P1 Transduction”. In: Current Protocols in Molecular Biology. Curr Protoc Mol Biol 2007. 10.1002/0471142727.mb0117s79.

[38] Sprouffske K, Wagner A. Growthcurver: an R package for obtaining interpretable metrics from microbial growth curves. BMC Bioinformatics 2016 17:1 2016;17:172-. 10.1186/S12859-016-1016-7.

[39] Nishida S, Duan G, Ohkama-Ohtsu N, Uraguchi S, Fujiwara T. Enhanced arsenic sensitivity with excess phytochelatin accumulation in shoots of a SULTR1;2 knockout mutant of Arabidopsis thaliana (L.) Heynh. Soil Sci Plant Nutr 2016;62:367–72. 10.1080/00380768.2016.1150790.

[40] PhD RM, Thangavel P, Dhankher OP, Long S. Separation and quantification of monothiols and phytochelatins from a wide variety of cell cultures and tissues of trees and other plants using high performance liquid chromatography. Journal of Chromatography A 1207: 72-83 2008;1207:72–83. 10.1016/j.chroma.2008.08.023.

