## Supplementary Files for "Functional Analysis of *cha* Genes Identifies ChaC as a Glutathione-Degrading Enzyme Rather Than a Sodium Transport Regulator"

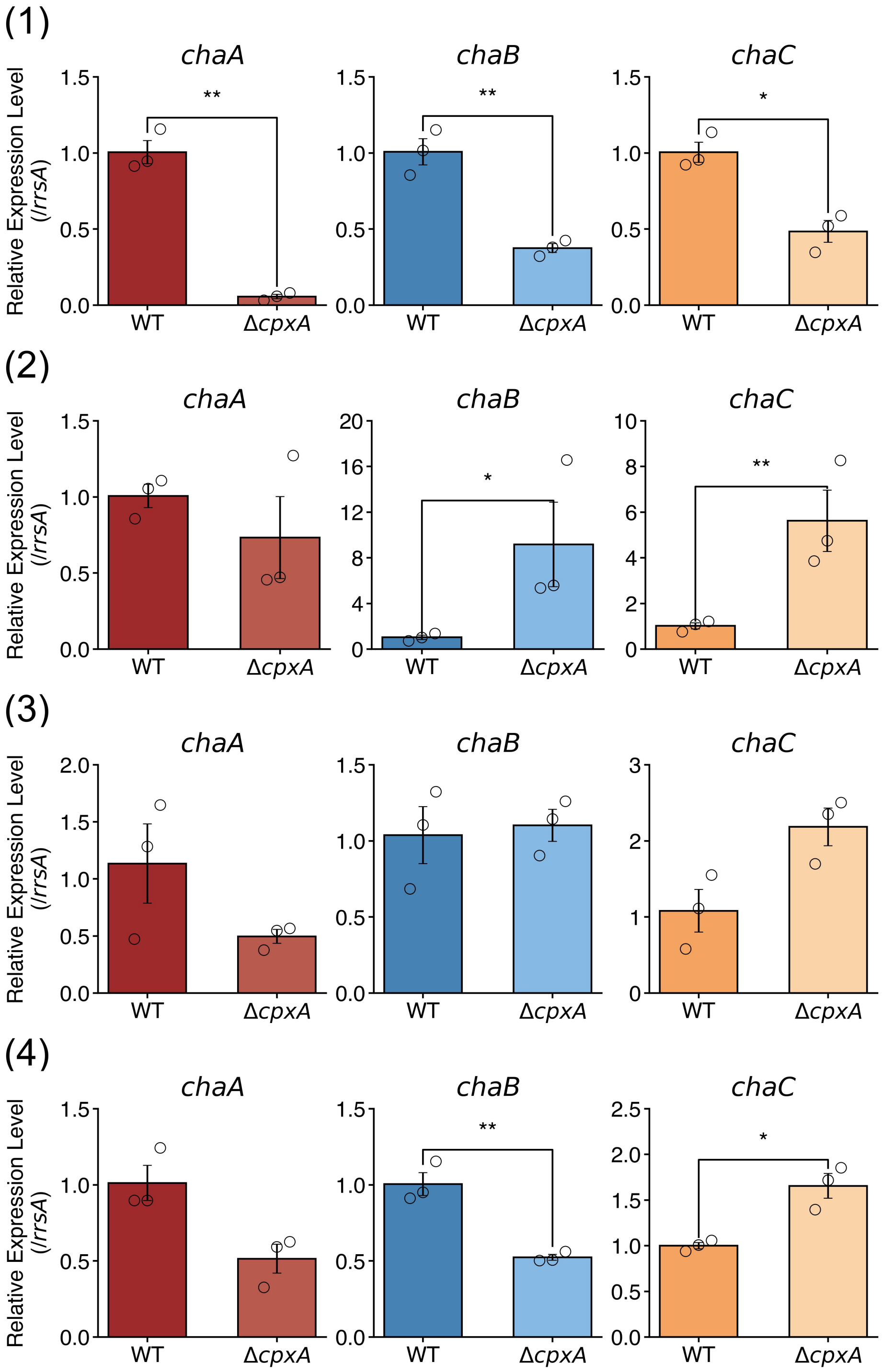


**Supplementary Figure 1. Transcript levels of *chaA*, *chaB*, and *chaC* in Δ*cpxA* mutant.** The values are presented as mean ± standard error of the mean (SEM) of $2^{\wedge-\Delta\Delta Ct}$ from three biological replicates. Numbers (1) – (4) represent four independent experiments. $\Delta C_T$ values were used for statistical analysis. Statistical analysis was performed using Student's t- test; *$P$ < 0.05, **$P$ < 0.01, ***$P$ < 0.001.

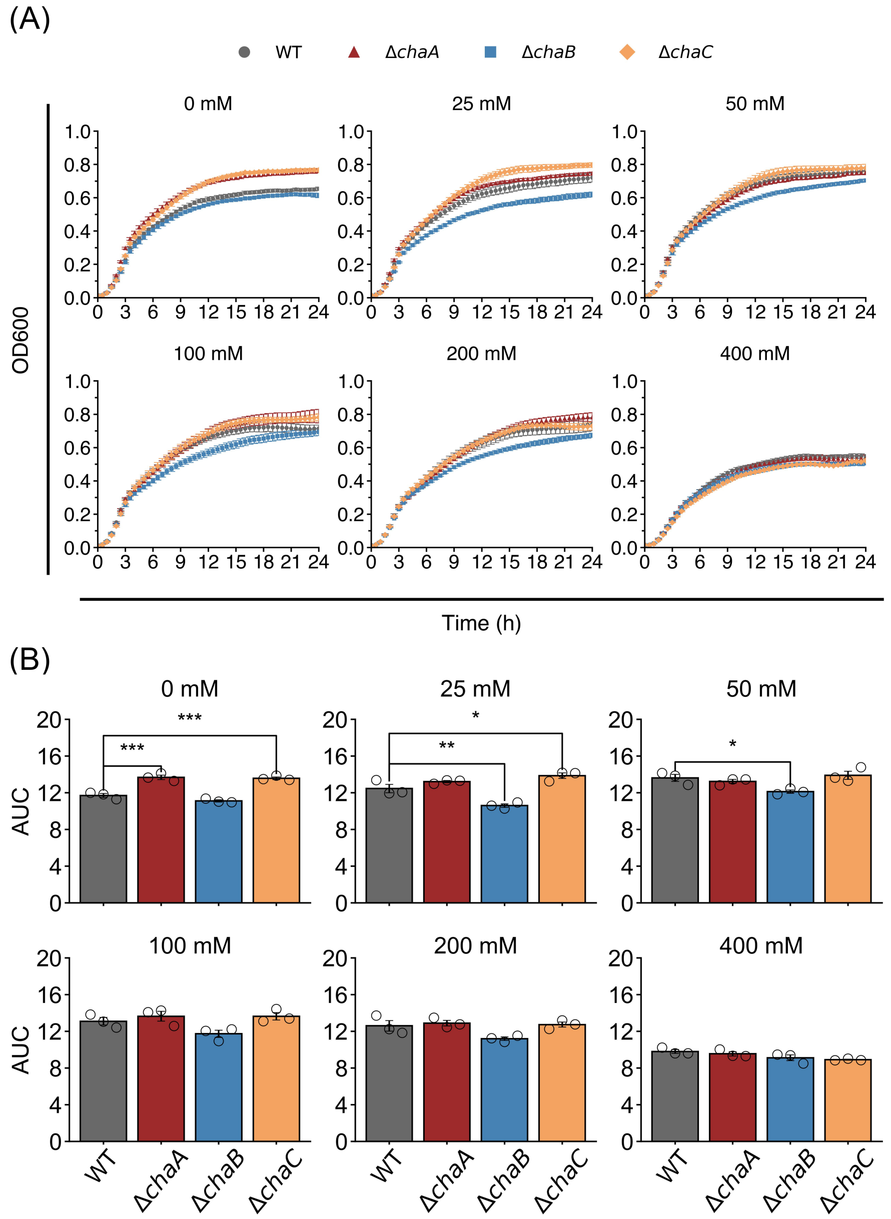


**Supplementary Figure 2. Effect of the *chaA, chaB*, or *chaC* deletion on salt sensitivity and growth in WT at pH 8.8.** (A) Growth curves of WT, Δ*chaA*, Δ*chaB*, and Δ*chaC* were generated by monitoring $OD_{600}$ from 0 to 24 h using a microplate reader during incubation in LBK medium containing 0, 25, 50, 100, 200, or 400 mM NaCl at pH 8.8 at 37 °C. Raw $OD_{600}$ values were corrected by subtracting blank measurements at each time point. (B) Area under the curve (AUC) values for WT, Δ*chaA*, Δ*chaB*, and Δ*chaC* were calculated from the growth curves shown in (A) over the 0 to 24 h interval using the R package *Growthcurver*. For panel (A) and (B), values are presented as mean ± SEM from three biological replicates. Statistical analyses were performed using one-way ANOVA followed by Dunnett's multiple-comparison test. *$P < 0.05$, **$P < 0.01$, ***$P < 0.001$.

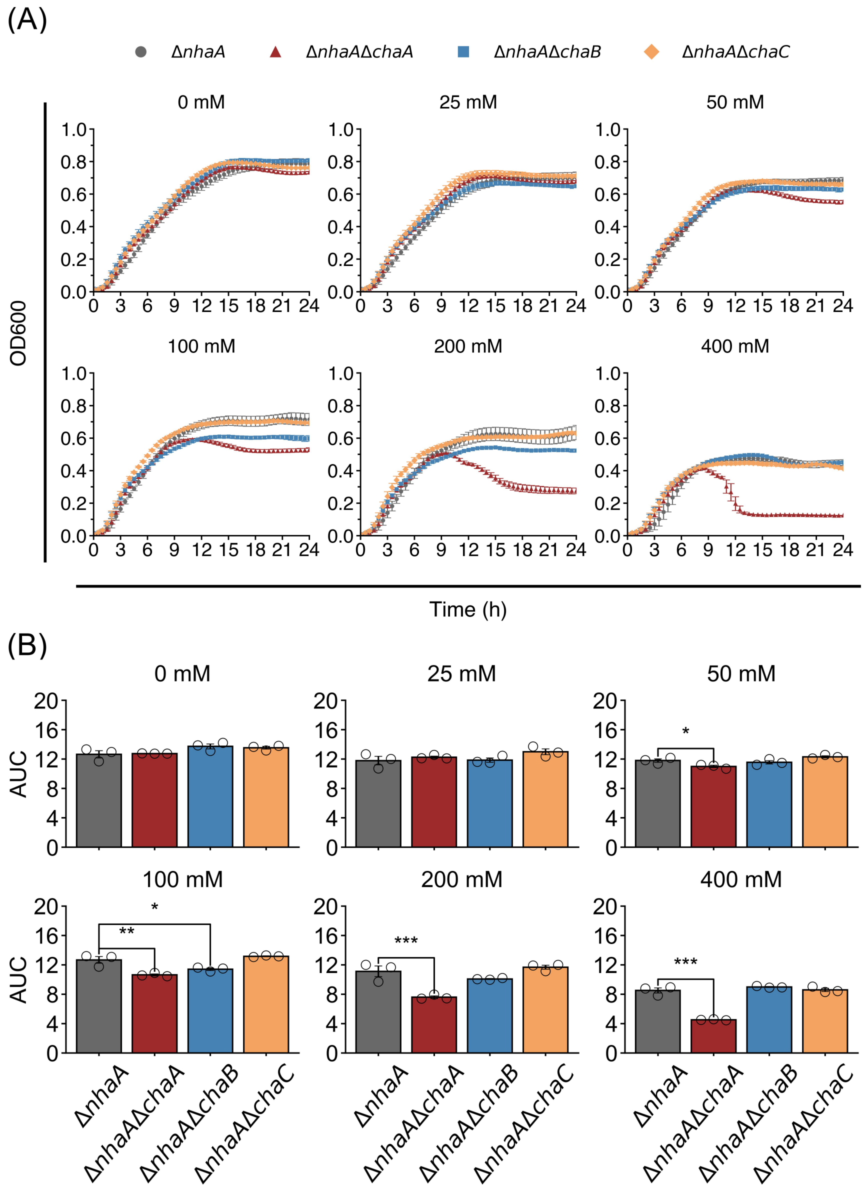


**Supplementary Figure 3. Effect of the *chaA, chaB*, or *chaC* deletion on salt sensitivity and growth in the Δ*nhaA* background at pH 7.0.** (A) Growth curves of Δ*nhaA*, Δ*nhaA*Δ*chaA*, Δ*nhaA*Δ*chaB*, and Δ*nhaA*Δ*chaC* were generated by monitoring $OD_{600}$ from 0 to 24 h using a microplate reader during incubation in LBK medium containing 0, 25, 50, 100, 200, or 400 mM NaCl at pH 7.0 at 37 °C. Raw $OD_{600}$ values were corrected by subtracting blank measurements at each time point. (B) Area under the curve (AUC) values for Δ*nhaA*, Δ*nhaA*Δ*chaA*, Δ*nhaA*Δ*chaB*, and Δ*nhaA*Δ*chaC* were calculated from the growth curves shown in (A) over the 0 to 24 h interval using the R package *Growthcurver*. For panel (A) and (B), values are presented as mean ± SEM from three biological replicates. Statistical analyses were performed using one-way ANOVA followed by Dunnett's multiple-comparison test. **P* < 0.05, ***P* < 0.01, ****P* < 0.001.

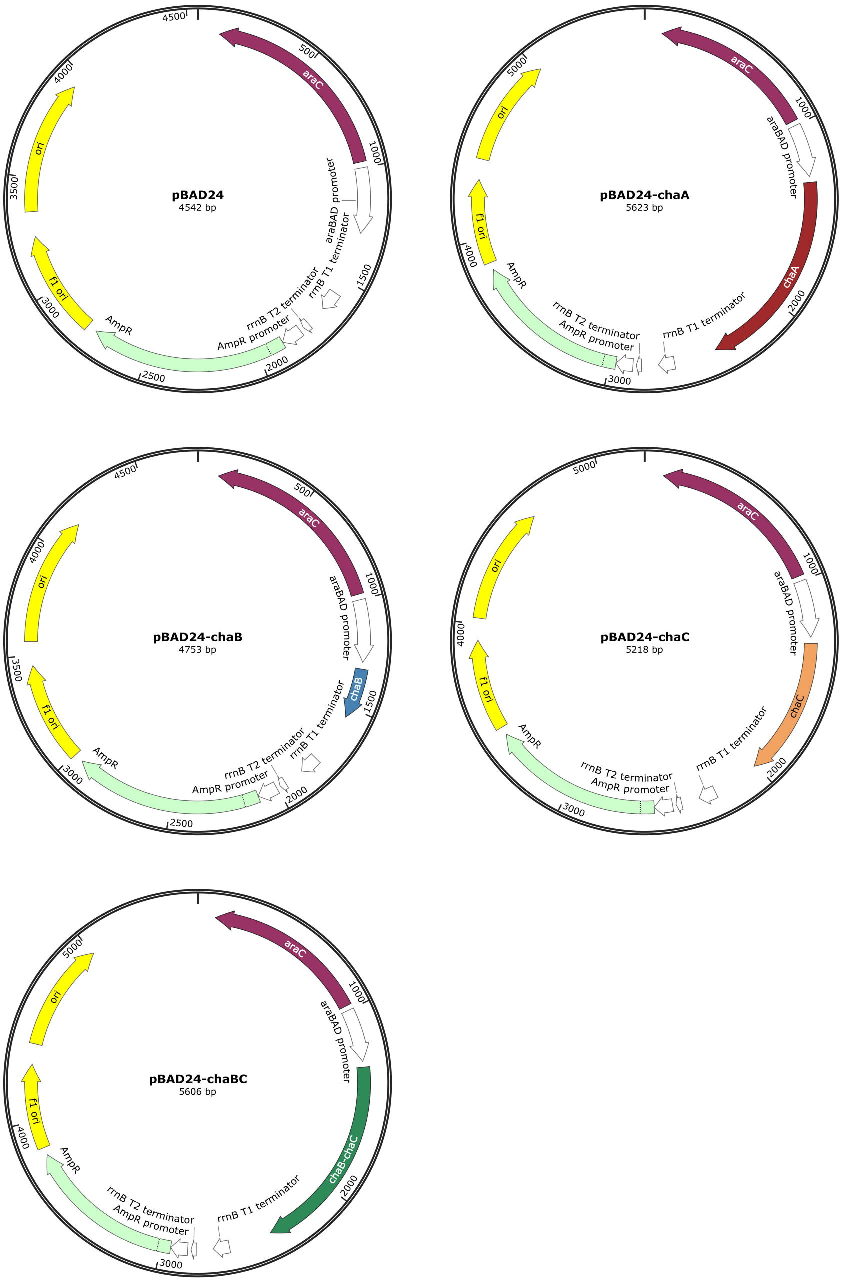


**Supplementary Figure 4. Maps of the plasmid for gene overexpression used in study.** The pBAD24 was obtained from National BioResource Project (NBRP). (A) pBAD24, vector control (EV), (B) pBAD24-chaA, for *chaA* overexpression, (C) pBAD24-chaB, for *chaB* overexpression, (D) pBAD24-chaC, for *chaC* overexpression, (E) pBAD24-chaBC, for *chaBC* overexpression. *chaBC* represents a contiguous genomic sequence spanning from the beginning of the *chaB* coding sequence to the end of the *chaC* coding sequence. The plasmid maps were generated using Snap Gene Viewer (from GSL Biotech; available at snapgene.com).

### Supplementary table 1. Bacteria strains used for this study

| Strain | Background | Plasmid | Antibiotic Resistance | Notes |
|---|---|---|---|---|
| W25113 (WT) | BW25113 | - | - | obtained from NBRP (NIG, Japan) |
| *chaA::kan* | BW25113 | - | Kanamycin | obtained from NBRP (NIG, Japan) |
| *chaB::kan* | BW25113 | - | Kanamycin | obtained from NBRP (NIG, Japan) |
| *chaC::kan* | BW25113 | - | Kanamycin | obtained from NBRP (NIG, Japan) |
| *nhaA::kan* | BW25113 | - | Kanamycin | obtained from NBRP (NIG, Japan) |
| *cpxA::kan* | BW25113 | - | Kanamycin | obtained from NBRP (NIG, Japan) |
| Δ*chaA* | BW25113 | - | - | *kan* was removed from *chaA::kan* |
| Δ*chaB* | BW25113 | - | - | *kan* was removed from *chaB::kan* |
| Δ*chaC* | BW25113 | - | - | *kan* was removed from *chaC::kan* |
| Δ*nhaA* | BW25113 | - | - | *kan* was removed from *nhaA::kan* |
| Δ*cpxA* | BW25113 | - | - | *kan* was removed from *cpxA::kan* |
| Δ*nhaA* Δ*chaA* | BW25113 | - | - | *nhaA::kan* was introduced into Δ*chaA* by P1 transduction, after which *kan* was removed. |
| Δ*nhaA* Δ*chaB* | BW25113 | - | - | *nhaA::kan* was introduced into Δ*chaB* by P1 transduction, after which *kan* was removed. |
| Δ*nhaA* Δ*chaC* | BW25113 | - | - | *nhaA::kan* was introduced into Δ*chaC* by P1 transduction, after which *kan* was removed. |
| EV (Empty Vector) | Δ*nhaA* | pBAD24 | Ampicillin | Δ*nhaA* transformed with pBAD24 empty vector |
| *chaA* ox | Δ*nhaA* | pBAD24-*chaA* | Ampicillin | Δ*nhaA* transformed with pBAD24-*chaA* |
| *chaB* ox | Δ*nhaA* | pBAD24-*chaB* | Ampicillin | Δ*nhaA* transformed with pBAD24-*chaB* |
| *chaC* ox | Δ*nhaA* | pBAD24-*chaC* | Ampicillin | Δ*nhaA* transformed with pBAD24-*chaC* |
| *chaBC* ox | Δ*nhaA* | pBAD24-*chaBC* | Ampicillin | Δ*nhaA* transformed with pBAD24-*chaBC* |
| DH5α | - | - | - | Preparation of plasmid DNA (Toyobo, Japan) |

#### Supplementary table 2. Primers used for this study

| Name | Sequence | note | Reference |
|---|---|---|---|
| chaA-F | CCAACGCTAATGCGTGATAC | For RT-qPCR analysis and mutant screening | |
| chaA-R | ATTGGGTGGCAAACTTACGA | For RT-qPCR analysis and mutant screening | |
| chaB-F | CACGTTCTACCGTCTCATGC | For RT-qPCR analysis and mutant screening | |
| chaB-R | CACTTTATGCGCGGTTTCTT | For RT-qPCR analysis and mutant screening | |
| chaC-F | GAATCCGGCACTGGAGTTTA | For RT-qPCR analysis and mutant screening | |
| chaC-R | CCCTCTTTCAGTGCAAGCAT | For RT-qPCR analysis and mutant screening | |
| cpxA-F | TGCGTGATGGCGAAGATAAT | For RT-qPCR analysis and mutant screening | |
| cpxA-R | GGCGTACTGACCAACATGG | For RT-qPCR analysis and mutant screening | |
| nhaA-F | GATTGTGCCGGCATTACTCT | For RT-qPCR analysis and mutant screening | |
| nhaA-R | GCCAGTACACCAAGTGCAAA | For RT-qPCR analysis and mutant screening | |
| rrsA-F | CTCTTGCCATCGGATGTGCCCA | For RT-qPCR analysis and mutant screening | Zhou et al., 2011 |
| rrsA-R | CCAGTGTGGCTGGTCATCCTCTCA | For RT-qPCR analysis and mutant screening | Zhou et al., 2011 |
| k2 | CGGTGCCCTGAATGAACTGC | For mutant screening | Datsenko and Wanner, 2000 |
| kt | CGGCCACAGTCGATGAATCC | For mutant screening | Datsenko and Wanner, 2000 |
| chaA-F2 | ATGTCAAATGCTCAAGAGGC | For RT-PCR analysis | |
| chaA-R2 | TCAGGCAAATATCGTCATCAA | For RT-PCR analysis | |
| chaB-F2 | ATGCCGTATAAAACGAAAAGC | For RT-PCR analysis | |
| chaB-R2 | TTACGATTTTTTATGCCATTTATCAT | For RT-PCR analysis | |
| chaC-F2 | GTGATAACGCGTGATTTCTTG | For RT-PCR analysis | |
| chaC-R2 | GTGATAACGCGTGATTTCTTG | For RT-PCR analysis | |
| chaA-F-EcoRI | AAAGAATTCATGTCAAATGCTCAAGAGGC | For insert amplification | |
| chaA-R-XbaI | AAATCTAGATCAGGCAAATATCGTCATCAA | For insert amplification | |
| chaB-F-EcoRI | AAAGAATTCATGCCGTATAAAACGAAAAGC | For insert amplification | |
| chaB-R-XbaI | AAATCTAGATTACGATTTTTTATGCCATTTATCAT | For insert amplification | |
| chaC-F-EcoRI | AAAGAATTCGTGATAACGCGTGATTTCTTG | For insert amplification | |
| chaC-R-XbaI | AAATCTAGATCAGGCGAATCCCGG | For insert amplification | |
| pBAD24-F | ATGCCATAGCATTTTTATCC | For insert verification | |
| pBAD24-R | GATTTAATCTGTATCAGG | For insert verification | |
